# A new framework for the study of apicomplexan diversity across environments

**DOI:** 10.1101/494880

**Authors:** Javier del Campo, Thierry Heger, Raquel Rodríguez-Martínez, Alexandra Z. Worden, Thomas A. Richards, Ramon Massana, Patrick J. Keeling

**Affiliations:** University of British Columbia, 3529-6270 University Boulevard, Vancouver, BC, V6T1Z4, Canada; Department of Marine Biology and Oceanography, Institut de Ciències del Mar (CSIC), Barcelona, Catalonia, Spain; Soil Science Group, CHANGINS, University of Applied Sciences and Arts Western Switzerland, Nyon, Switzerland; Living Systems Institute, Biosciences, College of Life and Environmental Sciences, University of Exeter, Stocker Road, Exeter, EX4 4QD, UK; Monterey Bay Aquarium Research Institute, Moss Landing, CA 95039, US

## Abstract

Apicomplexans are a group of microbial eukaryotes that contain some of the most well-studied parasites, including widespread intracellular pathogens of mammals such as *Toxoplasma* and *Plasmodium* (the agent of malaria), and emergent pathogens like *Cryptosporidium* and *Babesia*. Decades of research have illuminated the pathogenic mechanisms, molecular biology, and genomics of model apicomplexans, but we know surprisingly little about their diversity and distribution in natural environments. In this study we analyze the distribution of apicomplexans across a range of both host-associated and free-living environments, covering animal hosts from cnidarians to mammals, and ecosystems from soils to fresh and marine waters. Using publicly available small subunit (SSU) rRNA gene databases, high-throughput environmental sequencing (HTES) surveys such as Tara Oceans and VAMPS, as well as our own generated HTES data, we developed an apicomplexan reference database, which includes the largest apicomplexan SSU rRNA tree available to date and encompasses comprehensive sampling of this group and their closest relatives. This tree allowed us to identify and correct incongruences in the molecular identification of sequences, particularly within the hematozoans and the gregarines. Analyzing the diversity and distribution of apicomplexans in HTES studies with this curated reference database also showed a widespread, and quantitatively important, presence of apicomplexans across a variety of free-living environments. These data allow us to describe a remarkable molecular diversity of this group compared with our current knowledge, especially when compared with that identified from described apicomplexan species. This revision is most striking in marine environments, where potentially the most diverse apicomplexans apparently exist, but have not yet been formally recognized. The new database will be useful for both microbial ecology and epidemiological studies, and provide valuable reference for medical and veterinary diagnosis especially in cases of emerging, zoonotic, and cryptic infections.

**Author Summary:** Apicomplexans are important animal and human parasites, but little is known about their distribution and diversity in the natural environment. We have developed a phylogenetically informed and manually curated reference database for the SSU rRNA barcode gene, and analyzed all publicly available sequences from a broad range of environments, providing a needed framework to analyze high-throughput environmental sequencing (HTES) data. The reference database and the amplicon sequences identified a striking diversity of apicomplexans in the environment, including habitats not usually associated with this group (e.g. open ocean). The phylogenetic framework will underpin microbial ecology studies and provide valuable resources for medical and veterinary biology.

## Introduction

Protistan parasites account for more than 10% of the World Organization for Animal Health’s list of notifiable diseases of terrestrial and aquatic animals [1] and 25% of the major groups of pathogens that cause animal and plant species extinction and extirpation [2]. From an ecological perspective, it has been reported that parasites can represent a higher biomass than predators [3] and that they play key roles in the regulation and structure of natural communities in different environments [4]. Recent environmental microbial surveys also show that putatively parasitic protists are abundant in soils, freshwater, and marine systems [5–8].

Among these parasites, Apicomplexa stand out. Apicomplexans are obligate intracellular parasites that infect a wide variety of animals and are morphologically distinguished by the presence of an apical complex, a suite of structures used to invade host cells [9]. Most apicomplexans also possess a relict plastid, the apicoplast [10], which is non-photosynthetic but essential for parasite survival [11]. Well-known human parasites include *Toxoplasma* [12], *Cryptosporidium* [13], and the malaria agent *Plasmodium* [14]. Other apicomplexans are poorly studied, even though they are diverse and widespread in the environment and are hypothesized to play key roles in ecosystem function [7,8]. Of these, the gregarines are the largest but mostly understudied group of putative invertebrate parasites [15], also associated with juvenile frogs [16]. While the medical importance of apicomplexans is well-known, recent high-throughput environmental sequencing surveys (HTES) have shown a high diversity and abundance of apicomplexans in marine and terrestrial environments, suggesting our understanding of the impact of parasitic eukaryotes is probably underestimated. In the Tara Oceans global marine survey, apicomplexans represent the third most represented group of amplicons from parasites (following the Marine Alveolate groups I and II) [7]. In soils, apicomplexans are also highly represented in amplicon data, representing more than 50% of OTUs (Operational Taxonomic Units) in tropical soils [8].

Understanding what this diversity and distribution means requires a more detailed dissection of which apicomplexans appear in which environments. This is currently not possible because we lack a robust phylogenetic framework (e.g., a reference tree) upon which to base such inferences. Moreover, it has recently been shown that the apicomplexans are the sister group to another odd collection of microbial predators (colpodellids) and putatively symbiotic algae (chromerids), collectively known as chrompodellids or “Apicomplexan Related Lineages” (ARLs) [17–19]. These lineages have aided in understanding of how apicomplexans evolved to become parasites and the ecological conditions that might have led to this transition.

Most HTES studies use the small subunit (SSU) of the ribosomal RNA (18S rRNA for eukaryotes and 16S rRNA for prokaryotes) to identify/barcode the members of the targeted community. There are two major reference databases available for the 18S rRNA [20,21]. Both draw most of their taxonomic information directly from GenBank [22], which contains a large number of misidentified sequences, likely a historical and systematic problem derived from incomplete databases available when sequences are deposited. In one database, PR2, a significant number of groups have been curated by experts, but the phylogenetic framework for apicomplexans is under-developed, and in other databases, e.g. SILVA, there is minimal evaluation of the data as the data processing is automated [23,24]. The propagation of misidentified sequences often affects subsequent identifications, and ecological inferences more broadly, but in the case of pathogenic species (including apicomplexans) has also led to misdiagnosis of infections in humans [25].

Therefore, while we know from environmental survey data that apicomplexans and their relatives are widespread and diverse, no further interpretation is possible without a phylogenetically informed understanding of their sequence diversity. With that in mind, here we describe a comprehensive phylogenetic framework of all the apicomplexan and ARL sequence diversity, including those not identified, initially misidentified from all current isolates and environmental studies. We manually annotated the taxonomic information for each sequence using phylogenetic trees and curated associated information (such as the isolation source, origin, etc.) from the literature. As a product of this process, we propose several changes to apicomplexan classification, correcting the affiliation of several organisms and describing a dozen new groups. Using this reference framework as a tool, we examined the distribution of apicomplexans and ARLs in environmental data (both publicly available data and new data sets described here), covering environments from soils to the ocean, from sediments to the water column, at a level of taxonomic detail previously unachieved. Apicomplexans are shown to be present in all the environments examined, and more diverse than either previously thought or could be detected using current databases. Overall, these data represent a major tool for understanding the diversity and distribution of what are perhaps one of the most globally abundant animal parasites and highlights potential roles they might play in trophic networks in soils and marine systems.

These data also contribute to an overall understanding of the the parasitic nature of the group. Most available literature on apicomplexans define them as parasites. That has been shown for members of the better-known groups like coccicians or hematozoans, but has not formally been tested for most apicomplexans, especially the widespread and diverse gregarines. Most gregarines are nevertheless described as being parasitic because they were isolated from animal hosts, but in most of the cases Koch’s postulates have never been proven [26]. As large surveys based on molecular data become increasingly dominant in our view of microbial distribution, this problem becomes increasingly important because, for example, with HTES data we have fewer direct observations of the context of a host-microbe interaction. The level of apicomplexan diversity we document is incompatible with formal proofs of Koch’s postulates, further elevating the importance of clearly and accurately documenting patterns and correlations that will help to interpret the enormous amount of data generated from genomics and environmental genomics.

## Results

### A reference tree for apicomplexans

In order to improve our understanding of the molecular diversity of the apicomplexans, we constructed a 18S rRNA tree including all apicomplexan sequences retrieved from GenBank based on similarity and no shorter than 500 bp, resulting in a dataset which consists of 8,392 sequences representing a total of 756 distinct operational taxonomic units (OTUs) at 97% identity (Fig 1). From this curated phylogeny, we identified 12 novel environmental clades above the genus level within the apicomplexans, and 4 within the chrompodellids (Sup Info 1, Sup. Table 1). These clades consisted entirely of environmental sequences and included no known identified species. Additionally, the relationships identified between some well-studied groups also required alterations to the current taxonomic nomenclature. This was particularly clear for the genera *Babesia* and *Theileria*, where we identified five groups that had been identified as either *Babesia* or *Theileria,* but were not phylogenetically classifiable as being related to the type species. Both genera have minor SSU rRNA variants, but these tend to cluster with the major SSU rRNA variant from the same genus [27]. There is no indication our results result from undetected paralogues, and these OTUs should therefore be re-examined as they most likely represent new genera (Sup. Table 1). Similar situations were found for deep-branching eimerids and, less surprisingly, for gregarines, a group already known to be highly diverse and undersampled.

**Figure 1.**
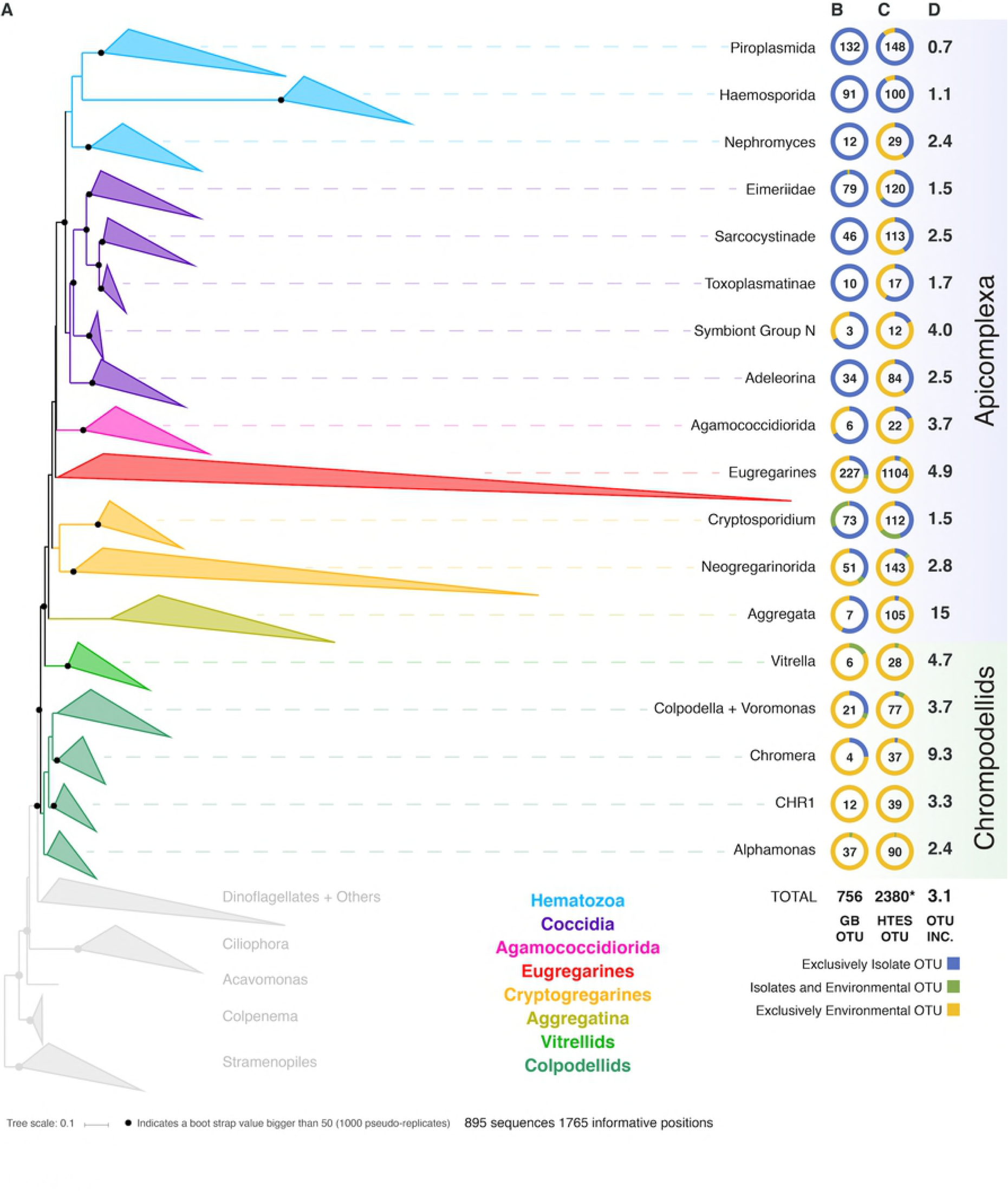
A) Apicomplexan reference tree inferred from a maximum likelihood (RAxML) phylogenetic tree constructed (best tree of 1000) using 18S rRNA sequences. Black dots represent bootstrap values above 50% (1000 bootstrap replicates). The tree has been collapsed in the main apicomplexan groups based on our taxonomic annotation (Supplementary Table 1). The tree showing all the OTUs is available as Supplementary Information 1. The first two columns next to the Apicomplexan groups names inform about the origin (colored circle) and the number of OTUs (number in the middle of the circle) retrieved from B) GenBank and from C) High-throughput environmental sequencing (HTES) studies. The HTES OTUs have been added to the reference tree using the Evolutionary Placement Algorithm in RAxML and the correspondent tree is available as Supplementary Information 2. The last column indicates the fold increase in numbers of OTUs after adding the HTES OTUs to the reference tree.

We then added all reads corresponding to apicomplexans and related lineages to this curated tree, incorporating more than 3.5 million sequences from HTES studies derived from diverse environments. We recovered a total of 2,380 apicomplexan related OTUs at 97% similarity (increasing the number of apicomplexan and ARL OTUs by a factor of three). The environmental sequences were not evenly distributed in the apicomplexan tree, but novel environmental diversity did appear in clades dominated by both clinical isolates and also groups that have been previously reported only in environmental surveys. In all cases, except the piroplasmids, the addition of HTES sequence data dramatically increased the proportion of OTUs retrieved from the environment compared with those from clinical isolates going from 270 to 1799 (Fig. 1). Overall, 30 new apicomplexan groups could be identified using HTES data, most of them within coccidians (Supplementary Table 2).

Along with the sequence data, we also curated all available metadata; either using information provided with the sequence accession, or by cross-referencing sequences with information in the publications describing the sequences. For clinical isolates this consisted of information such as host species or tissue type, while for environmental sequences it included location and the nature of the environment. Most of the publicly available sequences had associated metadata, and at the broadest level these corresponded to our expectations: most hematozoans and coccidians derived from isolates, while the rest of the apicomplexan groups were better represented in environmental surveys, particularly the eugregarines. In the case of the chrompodellids, most of the OTUs were also retrieved exclusively from environmental surveys, since there are relatively few cultures of these organisms.

### A reference database for apicomplexans and ARLs

We used the sequences from our comprehensive apicomplexan SSU tree to build a reference database that can be used to assign HTES reads an identity by sequence similarity and also to infer certain information regarding their environmental and host distribution. As for the tree the reference database consisted only of sequences longer than 500 bp (consistent with the previous analysis of 8,392 sequences). For each sequence in the database we provided an accession number and we manually assigned a phylogenetically-informed taxonomic string based on the tree that consists of six ranks below Eukarya and started with Alveolata. Right after the taxonomic string we included a column that corresponded to the name of the sequence. The name is either derived from a proper species name or a strain, but alternatively could also be a clone name in the case of sequences generated from environmental clone libraries. While the taxonomic string was assigned to each accession number based on the reference tree generated, the sequence names remained untouched for the purposes of identification within GenBank and the literature. Right after the taxonomic information we added the environmental metadata fields, starting with the origin of the sequence, if it came from an environmental clone library or from an isolate. After that, two columns with environmental information, named Environment 1 and Environment 2, with 1 being the most inclusive and 2 more detailed. More than 90% of the sequences had this field filled with 166 unique values for the less inclusive one. The next column was the geo-localization field, which was available for close to 90% of the sequences and populated by 715 unique values. The very last field was the host taxonomy string field. The string consisted of 5 taxonomic ranks, Phylum, Class, Order, Family and Species. Sixty four percent of the sequences contained information regarding the host taxonomy, using more than 700 unique terms at the species level. The metadata information was initially automatically retrieved from GenBank and after that double-checked with the literature that was also used to fill the significant gaps of the GenBank available metadata.

The metadata associated to the sequences retrieved from isolates was in most of the cases congruent with the information regarding tissue localization and host distribution of the correspondent organisms. When looking at the host distribution of the SSU rRNA sequences retrieved from GenBank (Fig. 2) we observed that in terms of distribution certain groups like the Agamoccidioria, the gregarines and Agregatta showed up mostly within the invertebrates, whereas coccidians, hematozoa, and cryptosporidium were obtained mostly from mammals. Considering the messiness of this kind of data in GenBank, it was a positive sign that at least for Apicomplexans this information could be reused and represented a good starting point to build a reliable database, especially if the information was double-checked and the gaps were filled using literature searches.

**Figure 2.**
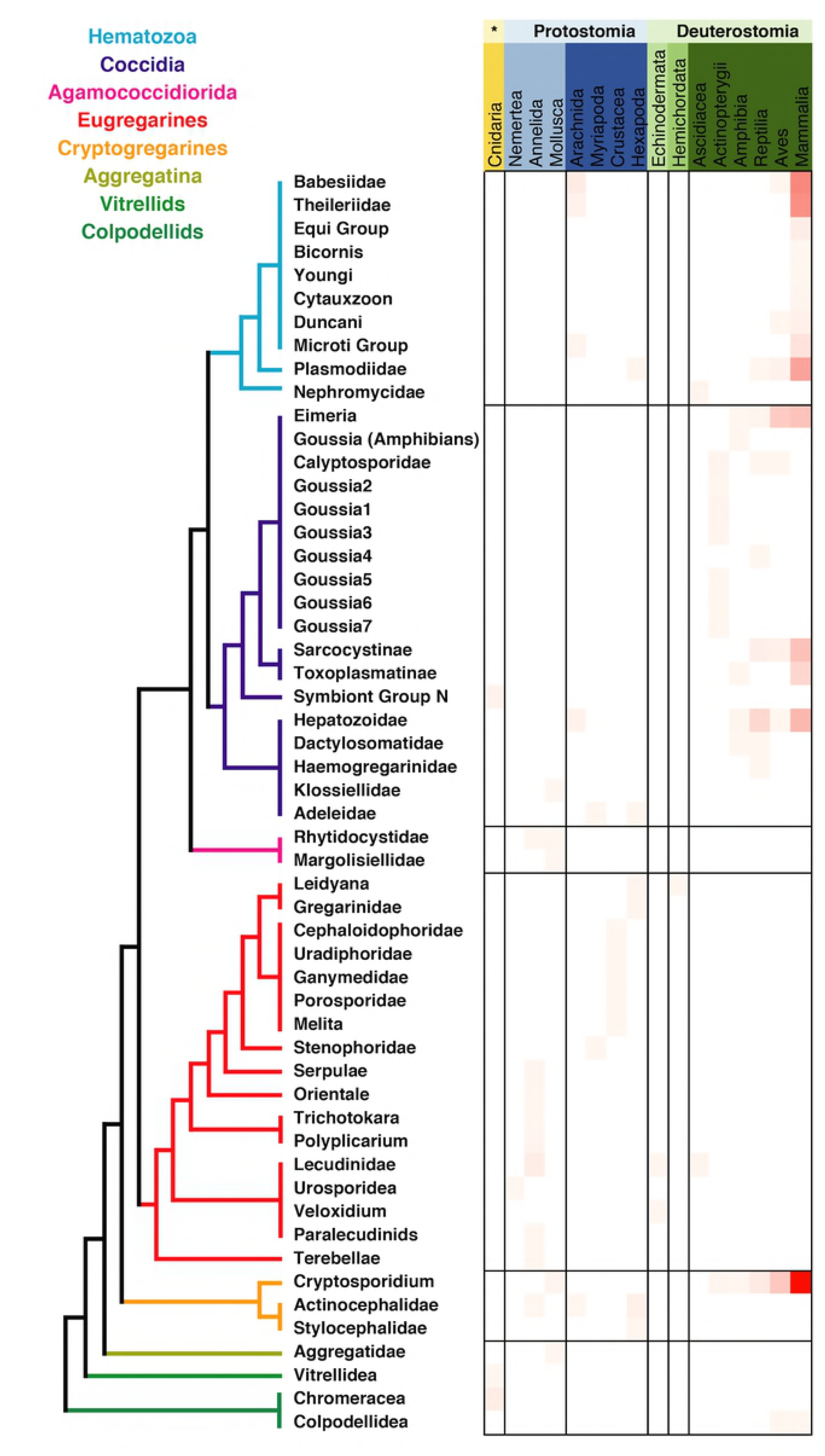
Apicomplexan host distribution heatmap based on the information associated to sequences available in GenBank.

### Environmental distribution

To evaluate the utility of the reference tree and database, and, at the same time study the environmental distribution of apicomplexans in existing environmental datasets, we analyzed 642 environmental surveys from 296 locations across different environments, including soil, freshwater, and marine habitats (Fig 3A, B). We retrieved apicomplexans from 76% of the samples and 94% of the environment-types, and found apicomplexans represented 0.6% of the amplicons as a whole. Overall, apicomplexans and their related lineage appeared to have a higher relative abundance in tropical than polar waters, and in marine and soil amplicon data as opposed to freshwater environments (Fig 3B). Comparing the apicomplexan sequences in HTES data with all available sequences in GenBank showed that the sequence similarity between retrieved reads and all other available sequences peaks around 93%, but when the comparison was made only against described species the peak drops to 84-85% (Fig 3C). This suggests that the vast majority of environmental sequences from apicomplexans come from yet-to-be identified species.

**Figure 3.**
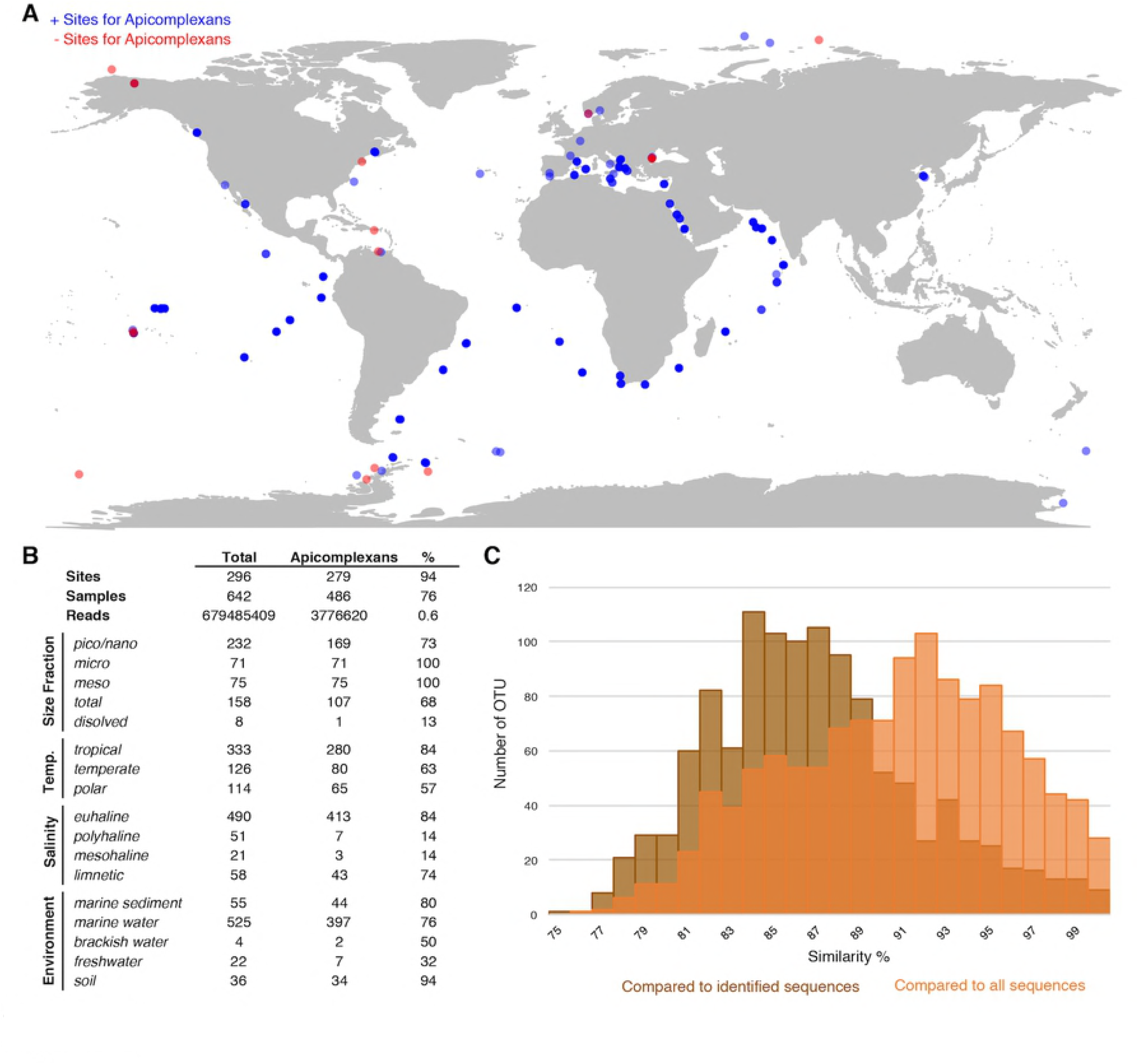
A) World map showing all the HTES analyzed sites, in red the sites were apicomplexan reads have been reported. B) Distribution (presence) of apicomplexans in the analyzed HTES datasets as a whole and clustered by different features. C) Blast Similarity against Genbank of the HTES apicomplexan reads. In brown the results when comparing against identified apicomplexans species and in orange the results when comparing against all the apicomplexan species including those environmental sequences that do not correspond to any known apicomplexan species.

One might expect environmental samples to yield mostly sequences associated with free-living clades, such as predatory chrompodellids, but we found the opposite. Most of the reads identified as apicomplexans from environmental surveys fell within the eugregarines, which is perhaps not surprising since they are known to be diverse and are less sampled than other apicomplexan groups. The gregarine life cycle also involves releasing large numbers of resistant, infective cells into the environment, rather than direct, host-to-host infection, which might also be expected to lead to high relative abundance representation in environmental data. Eugregarine amplicon abundance was closely followed by that of the comparatively well-studied coccidians, where a substantial environmental diversity of adeleorinids, sarcocystids and basal *Goosia*-like eimerids, and agreggadids was identified (Fig 4). We also retrieved sequences corresponding to cryptogregarines and more surprisingly, hematozoans. Sequences belonging to the chromopodellids were also identified, including both members of clades containing free-living genera like *Alphamonas, Voromonas*, or *Colpodella*, as well as genera thought to be symbionts such as *Chromera* and *Vitrella*. There were no significant differences between the distribution of 18S rRNA V4 and V9 reads across the tree that cannot be explained by the source of the sequences.

**Figure 4.**
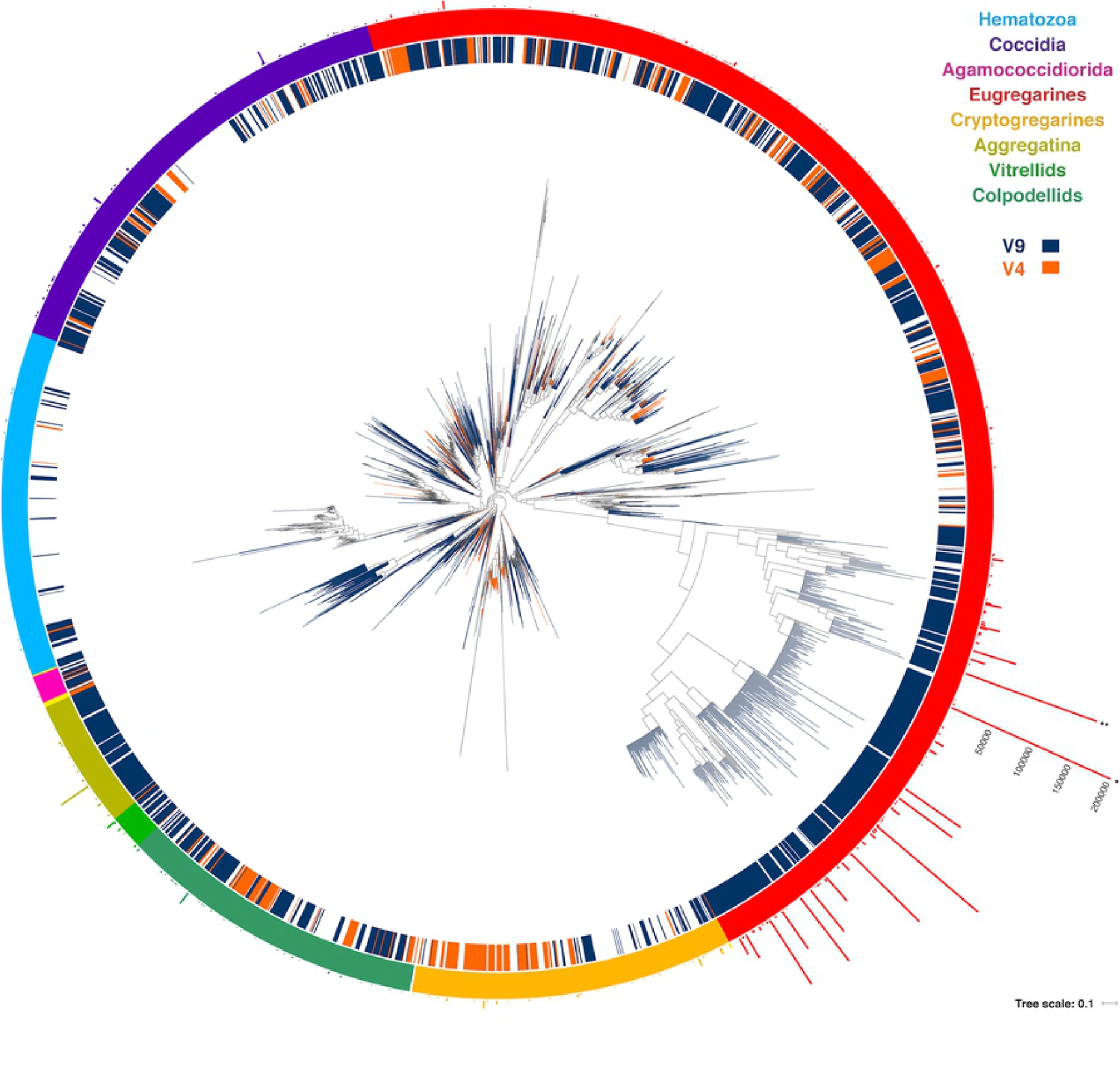
Round tree showing the placement of the short reads using the Evolutionary Placement Algorithm in RAxML (EPA-RAxML) on the reference tree. The colors of the leaves and the inner crown surrounding the tree indicate the 18S rRNA region of the corresponding OTUs (V4 or V9). The outer crown indicates the number of reads per OTU.

When comparing the environmental distribution using both Sanger and HTES sequences (Fig 5), we observed that hematozoans and coccidians were commonly retrieved from clinical isolates but rarely observed in Sanger environmental clone libraries. However, they did appear in marine, freshwaters and soils HTES datasets. We retrieved OTUs from well-known groups but also from 11 novel clusters that could not be assigned to any of the described hematozoans or coccidians. In the case of the eugregarines and cryptogregarines, the increase in diversity from HTES data was extreme: members of both groups have been retrieved from isolates in the past, but because nearly all inhabit invertebrate cells relatively few have been previously characterized. The addition of HTES data confirmed the abundant representation of eugregarines and cryptogregarines in the environment, and showed that they had the highest relative abundances among amplicons along with being the most widespread and diverse.

**Figure 5.**
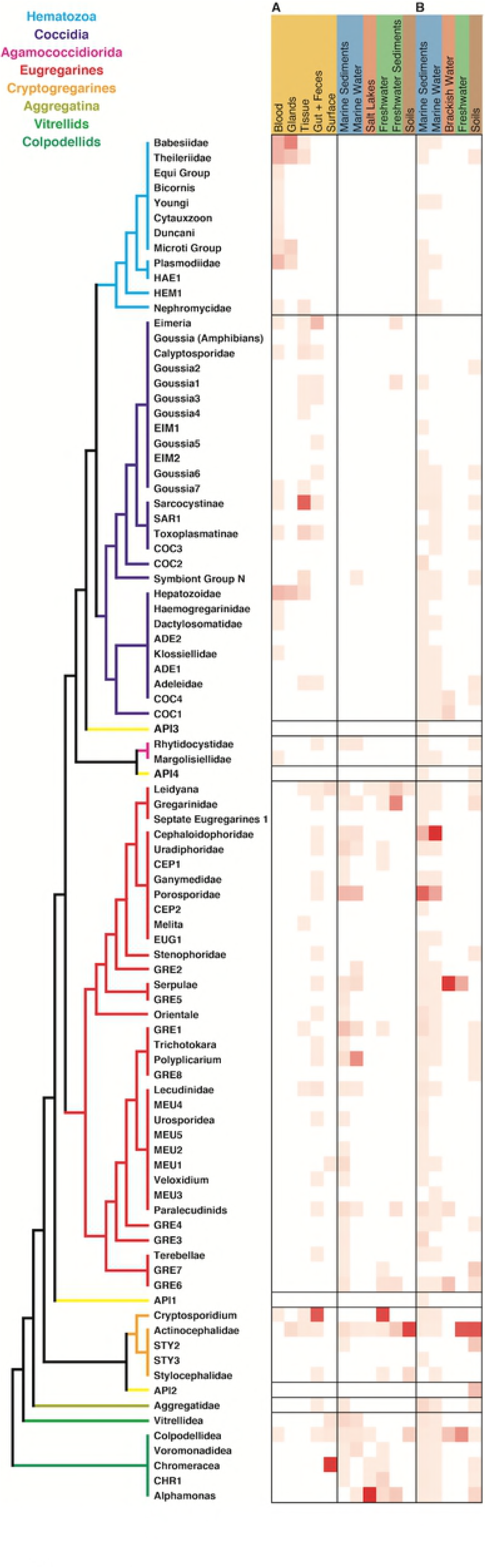
Apicomplexan environment distribution heatmap based on the information associated to the sequences retrieved from A) GenBank and B) relative amplicon abundances in HTES studies. The novel lineages obtained only by HTES studies are inserted on the tree shown in Fig. 2, respecting the original topology.

In marine systems the Cephaloidopphoridea and Porpospodidae eugregarines had the highest relative abundances among apicomplexans, while in freshwaters and soils the Actinocephalidea cryptogregarines had the highest relative abundances. In the marine environment, the most highly-represented groups in open ocean were the Cepholoidophoridae and Porosporidae gregarines, while in coastal environments the Lecudinidae, GRE1, and Chromareraceae dominated the apicomplexan derived amplicons (Sup Fig 1). Cepholoidophoridae and Porosporidae were the most highly-represented in epipelagic and benthic environments, while Dactylosomatidae, Klossiellidae, Adeleidae, and Gregarinidae were the most common in mesopelagic and bathy/abyssopelagic environments. Apicomplexans are typically aerobes, and all of the groups were found from oxic environments, but a few were also identified in anoxic ones, where Dactylosomatidae, Klossiellidae, and Gregarinidae showed the highest amplicon abundances.

## Discussion

### A framework for biomedical and ecological studies

The widespread use of HTES data to infer characteristics about the distribution and ecology of microbial life relies entirely on the quality of the reference database to translate the catalogue of sequences recovered from an environment into accurate taxonomic identifications of the organism from which they are derived. Despite their importance, available databases have not been phylogenetically curated for most protist lineages. A common practice is instead to export annotations from GenBank, where many sequences are mis-annotated or not annotated at all. As a result, GenBank is the de facto reference database for users who may be unaware of how to access specific references databases.

In the case of the apicomplexans, the problems with the current state of our reference data are clear from comparisons of our manually curated reference database and tree with existing GenBank annotation. There are extensive mistakes in some human and animal parasites, like *Theileria* and *Babesia* [28] at the genus level (Sup Table 1), or in *Cryptosporidium* and eimeriids at the species level (Sup Table 1). Mis-annotation is even more common in the eugregarines and the cryptogregarines at various taxonomic levels, and mistaken affiliations are relatively common across the entire tree of apicomplexans (Sup Table 1). Putting this in a biomedical or veterinarian context, an incorrect annotation of a genetic barcode in a reference database can lead to mis-diagnosis and potentially ineffective treatments. For ecological studies the situation is much worse, because the environmental sequences in GenBank are usually not annotated taxonomically at all, so even a perfect sequence match leads to no information about identity. To give an example of how this leads to problems, if one looks at the protists reported in the Tara Oceans survey of marine microbes, they describe the widespread presence of the malaria parasite in marine samples [7]! Specifically, the Tara Oceans analysis recovered 1,123 OTUs representing 293,824 reads out of 6 million aplicomplexan reads that were widespread across the photic zone, and these were assigned as *Plasmodium falciparum*. Analyzing the same data using our phylogenetically curated reference framework, we have retrieved only two OTUs representing a total of nine Tara Oceans reads that are phylogenetically affiliated with *Plasmodium*: this is three orders of magnitude fewer *Plasmodium* OTUs and five orders of magnitude fewer reads (Sup Table 2). The large number of “*Plasmodium*” reads that were mis-identified using existing reference databases are in fact mostly Cephaloidophoridea, gregarine parasites of crustaceans (see below): the ecological implications of this mis-identification needs no elaboration.

### Diversity and distribution of clinically important apicomplexans

Based on the publicly available information in GenBank, one would conclude that the most common apicomplexans are human- or cow-parasites, such as *Plasmodium, Babesia*, or *Cryptosporidium* (Fig 2, Sup Table 1). However, this is an obvious reflection of the biases in apicomplexan research foci, and GenBank offers little to aid our understanding of lineage distributions. The biological diversity of apicomplexans as inferred from HTES data is dominated by gregarines, particularly the eugregarines, of which nearly all known species inhabit invertebrates. The biases in GenBank are even more prominent when we examine the similarity between reference sequences and HTES sequences, which on average share only 85% similarity to identified species. This value is extremely low, and indicates that not only has the majority of apicomplexan diversity not been characterized at the molecular level, but that we lack even representatives of many major lineages for comparison. A large-scale screen of invertebrate-associated apicomplexans would greatly help to improve our knowledge about the diversity, the ecology, and the evolution of the apicomplexans, because we have barely begun to scratch the surface of this part of the apicomplexan tree.

Within the cultured apicomplexans, there is an obvious bias in favor of clinically important species as well. Among these clinical isolates, blood borne pathogens from mammals (and particularly from bovids and humans) are dominant, and there are also significant numbers of *Cryptosporidium* isolates from guts and faeces: all reflecting the interest of the medical and veterinary communities. Interestingly, most of these well-known pathogens (with the exception of *Cryptosporidium*) are not retrieved from Sanger clone library environmental sequence surveys, but they do appear in HTES data, especially from soils and marine samples. Nevertheless, the phylogenies of well-known pathogens are significantly improved by analyzing the data as a whole, including all the other apicomplexans and both sequences from isolates and from environmental surveys. As mentioned above, *Babesia* and *Theileria* are paraphyletic, suggesting the need for a taxonomic revision of these groups. Similarly, *Cryptosporidium* isolates are taxonomically problematic at the species level, and at some level so is *Toxoplasma*.

The presence of sequences from important animal- and human-parasites in the free-living environment can also help us to identify unsuspected reservoirs, or even novel parasite diversity. Biomedical research tends to focus on strains already important to human disease, not unreasonably, but it would also be useful to examine HTES datasets more carefully to identify the natural distribution of apicomplexan lineages of clinical and veterinary interest outside these hosts, since these might reveal potentially undescribed sources of parasite transmission and propagation.

### The environmental distribution of the apicomplexans and related lineages

Apicomplexans appear in all environments examined, as has already been suggested by previous analyses [7,8]. There appears to be a bias towards marine and soil samples, but this may be due to a bias in the number of samples coming from these environments as brackish or freshwater environments are relatively under-studied (Fig 3), and they are not rare in samples that do exists from such environments. Their broad distribution and high abundance both suggest apicomplexans play a significant role in food webs and the population structure of their animal hosts.

When looking more closely at the distribution by group, the relative amplicons abundances of eugregarines suggest that they are the most abundant apicomplexans, that they are the most diverse (at least for the variable regions of the 18S rRNA gene that have been studied), and also have the widest spatial distributions (Figs 4, 5). The neogregarines are particularly well-represented among apicomplexan amplicons in soils, but not as diverse as eugregarines. Other groups that stand out are the adeleorinids, the eimeriids, and the sarcocystids, which are mostly present in marine samples. Adeleorinids includes some described genera that are extremely highly-represented, like *Klossia*, which infects mollusks [29], or fish-parasites in the Dactylosomatidae [30], which are the most prevalent apicomplexan amplicons in the deep ocean and also anoxic environments (Sup Fig 1). In the case of the eimeriids, most of the sequences fall at the base of the group in a paraphyletic assemblage including the fish-parasite *Goussia* [31]. Fish-parasites have also been described among sarcoystids and other coccidians [32,33], but little is known about their biology or diversity, and virtually nothing from a molecular perspective. Most of the coccidian sequences come from the sediment, and based on their known host distribution it seems plausible that this signal corresponds to the spores of fish parasites [34].

In other cases, observed distribution patterns are more surprising, in particular, the apicomplexan known as “Symbiont N”, and the apicomplexan-related genera *Vitrella* and *Chromera*. All these groups have been described as coral symbionts [35] or associated to coral reefs [36] and, although we do not have samples from any coral environment in our dataset, sequences from all three were retrieved in our analyses. Finding such sequences from sediments may simply raise the possibility that these organisms associate with other anthozoans and not just corals, but we also identified them in water column samples. This may be due to capturing infective stages between hosts, may suggest that these organisms are not necessarily symbionts, or that closely-related sisters are free-living. Most strangely, however, reads associated with all three groups of coral symbionts were also retrieved from soils. This is much harder to rationalize with their being coral symbionts and indicates that some of the biological diversity of these groups has not yet been explored.

A similar situation is surprisingly seen in the hematozoa. Hematozoans are best known for infecting terrestrial animals, but 2,483 hematozoan amplicons (Sup Table 2) were retrieved from marine samples, predominantly sediments (Fig 5). Half of these belong to the ascidian symbiont, *Nephromyces*, which is not unexpected since this is a marine host, but most other reads were closely-related to well-known blood borne parasites best known from non-marine hosts, like *Babesia* or *Plasmodium*. Neither Haemosporidia nor Piroplasmida are not known to infect fish, and this together with the relatively low representativeness of these reads suggests alternative origins such as marine birds, where these parasites have been previously reported [37].

### Gregarines as the most abundant apicomplexans and their putative roles in the environment

Eugregarines and neograrines are the most highly represented apicomplexans in amplicon data from marine and soil samples, respectively. This is not surprising because they are largely invertebrate-associated, and invertebrates represent the majority of animals, both taxonomically and numerically [38]. These two groups of apicomplexans are poorly-studied, but it seems likely that some of the more common but unidentified gregarines are associated with zooplanktonic and meiofaunal animals, which play crucial roles in the food webs (and so too, by extension, would their associated microeukaryotes). The role of gregarines as pathogens or parasites is still debated and there is every reason to expect some variation in their relationship to their hosts. Gregarines are known to cause disease in shrimps [39] and in insects they are involved in the decrease of their host fitness by tissue damage, reducing their body size, fecundity and longevity [40–42]. In most species, however, their pathogenicity has not been examined specifically.

The most abundant apicomplexan group in our dataset is the Cephaloidosporidea, which also represent one of the most abundant OTUs in Tara Oceans (where, as noted above, they are mis-identified as *Plasmodium*) [7]. Cephaloidophoroidea gregarines infect crustaceans from both Mallacostraca and Maxillopoda [38, 43] (Fig 2), and are known to infect copepods [44]. Considering the size fractions in our marine water samples (from 0,8 to 2000 µm), and the reported abundance of Cephaloidophoroidea in previous publications [7], it seems likely that they infect marine zooplankton in large numbers. Another group of eugregarines, the Porosporidae, also infect crustaceans, including copepods [43]. This group of eugregarines are phylogenetically related to the Cephaloidophoridea and were also commonly retrieved from both the water column and sediments. Both these groups may therefore play a major role in the marine food webs by regulating copepods populations, which are themselves the most prominent members of the zooplankton and a key link between phytoplankton and fish larvae [45]. Examining the co-occurrence of these gregarines with members the zooplankton could allow this hypothesis to be tested, while screening copepod individuals in the wild would be needed to conclusively confirm whether they are infected with these common but unidentified Cephaloidophoridea and Porosporidae, and whether the apicomplexans cause disease or death of the hosts.

In soils, the neogregarines are the most abundantly represented group in apicomplexan amplicon data, with the Actinocephalidae standing out. Related organisms are commonly retrieved from bees [46], fleas [47], and earthworms [48]. Apart from the difficulty of defining what is and what is not a pathogen, there are also infections that we do not know which disease they cause, if any. In the case of earthworms, *Monocystis* is an extremely common apicomplexan, with 100% infection rates in certain communities [48]. Earthworms play a significant role in the soil food web, where they are responsible of organic-matter breakdown, nutrient enrichment, particles relocation, and the dispersal of microorganisms, altogether shaping the soil structure and physic-chemical properties. In the case of *Monocystis*, there is no clear evidence it has a significant ecological impact, but it has been shown to be mildly deleterious to host fitness [48], so could only play a subtler role in host population structure, which in turn could be significant. Apicomplexans more broadly have been retrieved in high densities in tropical soils, with eugregarines and neogregarines dominating [8]. As with the ocean, some of these gregarines must be relevant to regulating host population diversity and activities that affect soil structure and composition. Such conclusions await concrete identification of apicomplexan-host associations as well as determining whether infection leads to disease or death, and the database has provided the first step of identifying which parasites to focus on.

Apart from a putative role as host-regulators, the wide distribution and high representativity of apicomplexans suggests they may also represent an alternative heterotrophic pathway for transferring carbon within the trophic web. Symbionts’ consumption rates are high in the environment, and this consumption is frequent, non-accidental, and influences food web properties [49]. On the other hand, when parasites infect their hosts they have access to more organic matter than free-living heterotrophic species that depend on prey encounters [50]. Overall, apicomplexans, and eugregarines and neogregarines in particular, might make a significant impact on food web dynamics and the carbon cycle in marine and soil systems simply through their heterotrophic activities, and not just on how they change host population numbers and structure.

## Conclusions

Gregarines (eugregarines and neogregarines) were identified as the most abundantly-represented and widespread apicomplexans in our analyses. Considering that previous studies have shown that the apicomplexans are also well-represented at the amplicon level compared with the rest of the micro-eukaryotes, gregarines as putative invertebrate parasites have the potential to play an important role regulating the meiofaunal and zooplanktonic communities in soil and marine systems, directly impacting the carbon cycle. It will be important to determine exactly the functional role that gregarines play in these environments, and its impact, by examining patterns of host-parasite population change together with more direct observations of their interactions. Only then can apicomplexans be realistically integrated into models for marine and soil trophic networks.

HTES metabarcoding has been an extremely useful tool for microbial ecology, and recently eDNA metabarcoding is becoming standard for rapid screening of organismal diversity in conservation [51] and environmental monitoring [52], providing a useful tool for diagnosis. However, it is crucial to have reliable reference databases to accurately identify the sequences generated through HTES. Our study provides the necessary tools to study the diversity and ecological distribution of the apicomplexans and establishes the basis to use the 18S rRNA gene as a reliable biomarker to detect apicomplexans in host associated and free-living environments, and by extension its use in epidemiology or diagnosis. Having this reference framework, consisting on a tree and a reference database, is the only way to interpret such data in a useful and comparable way.

We have shown the suitability of the aforementioned approach and tools by analyzing a large quantity of HTES data from public sources and generated *de novo*. Using the described framework, we have shown that apicomplexans are diverse and widespread based on their amplicon distributions in the environment. We cannot directly infer their organismal abundance using our dataset, but based on previous publications [7,8] they are one of the most highly-represented parasites in terms of 18S rRNA amplicon relative abundances, particularly in soils and marine systems. The novel diversity revealed here includes unrecognized parasites of humans and a range of ecologically and commercially important animals, in addition to several potential emergent pathogens. From a veterinary and medical perspective, it would be interesting to use eDNA techniques in the future to explore the prevalence of apicomplexans in the environment and target potential sources of infection.

## Materials and Methods

### Sample processing and sequence generation

Soils samples were obtained from Calvert and Hecate islands on the central coast of British Columbia, Canada. Samples were stored in coolers containing ice packs and afterward frozen at −80°C within 6 hours. DNA extraction of the samples was performed with the FastDNA SPIN Kit for soil (MP Biomedicals, Solon, OH, USA). Protist communities were investigated using high-throughput Illumina sequencing on the hypervariable V4 region of the 18S rRNA gene using using the Phusion^®^ High-Fidelity DNA Polymerase (Thermo Fisher, MA, USA) and the general eukaryotic primer pair TAReuk454FWD1 and TAReukREV3 [53]. Paired-end sequencing of the library was performed with the Illumina MiSeq platform using the MiSeq Reagent v3 chemistry (Illumina, San Diego, CA, USA). The library was 300bp paired-end sequenced at the Genotyping Core Facility of the University California Los Angeles (Los Angeles, CA, USA). Further details on sampling and sequence generation can be found at Heger et al. 2018 [54]. Amplicon data are available on NCBI Sequence Read Archive (SRA) under project number: PRJNA396681.

Marine sediment samples were taken on-board of the MBARI research vessel *Western Flyer* in the North Pacific Ocean (Monterey Canyon) and preserved with RNA Lifeguard (Qiagen, CA, USA). From each sampling core ∼12 g of sediment were transferred into a 50 mL falcon tube, using sterilized spatula in laminar flow hood. Samples were placed immediately in a –80 °C freezer on board. RNA extraction was performed using the Qiagen PowerSoil RNA isolation kit (Qiagen, CA, USA), using the DNase treatment described in the protocol. RNA quality and quantity for samples was checked using a 2100 Bioanalyzer (Agilent technologies). Each RNA sample was then reverse transcribed into cDNA using SuperScript III reverse transcriptase (Invitrogen, CA, USA) with random hexamers. Respective negative controls were done during the process. Protist communities were investigated using high-throughput Illumina sequencing on the hypervariable V9 region of the 18S rRNA gene using the Phusion^®^ High-Fidelity DNA Polymerase (Thermo Fisher, MA, USA) and the general eukaryotic primer pair 1380F and 1510R [55]. Paired-end sequencing of the library was performed with the Illumina HiSeq 2000 platform (Illumina, San Diego, CA, USA) with NEXTflex DNA sequencing kits and an identifying NEXTflex DNA barcode with 8-base indices (Bioo Scientific, TX, USA). The library was 150bp paired-end sequenced at the Exeter Sequencing Service (University of Exeter, UK). Amplicon data are available on NCBI Sequence Read Archive (SRA) under project number: XXXXXXX.

### Reference phylogenetic tree and database

All GenBank SSU rDNA sequences identified as Apicomplexans or Chrompodellids were retrieved using the corresponding taxid (5794 / 877183 & 177937 & 333132). Sequences shorter than 500 bp were excluded. The remaining sequences were clustered at 97% identity using USEARCH v7.0.1090 [56]. In order to build the tree, 22 other alveolates and 18 sequences were used as outgroups. All sequences were aligned and trimmed using MAFFT 7 with default settings [57] and trimAl [58], respectively. A maximum likelihood phylogenetic tree was constructed with RAxML 8.1.3 [59] using the rapid hill climbing algorithm and GTRCAT evolutionary model. Whether sequences belong to the Apicomplexans and the Chrompodellids was determined based on the tree topology and literature. Verified sequences were then used to iteratively retrieve more sequences from GenBank using blastn [60] against nt as previously described [61,62] in order to enrich the tree with environmental sequences or sequences with a wrong taxid that were not recovered in the first place. Putative chimeric sequences were manually examined and the final retrieved dataset was clustered as well at 97% in order to build a tree. The final phylogenetic tree was built using RAxML with the settings mentioned above. Statistical support for the consensus tree was calculated using non-parametric bootstrapping with 1,000 replicates.

In order to construct a reference database sequences from isolates were initially annotated based on previously published works. We adopted the established taxonomy as our default classification method when possible. For groups containing isolates with no formal taxonomic affiliation assignable based on the tree an informal name for the group has been provided based on the genus, species name of associated metadata. In the case of groups containing only environmental representatives a group named using three letters and a number has been provided. Metadata for the sequences in our dataset was downloaded from GenBank using custom scripts. For sequences still missing environmental data, their information was then collected manually from the literature.

### Analysis of HTES sequences

Sequences annotated as Alveolates were retrieved from three publicly available 18S rRNA datasets, VAMPS, BioMarKs and Tara Oceans [7,63,64] and two additional datasets generated by us. Overall the analyzed data covers a wide range of environments from soils and freshwater to the sunlit ocean and the deep-sea sediments. The analyzed dataset contains both V4 and V9 region reads and several size fractions. The fasta file containing all reads was sorted by length using USEARCH and clustered into OTUs with 97% similarity using QIIME with default setting (UCLUST). OTUs were then aligned with the reference alignment using PyNAST [65] embedded in QIIME [66] (align_seqs.py). The reference alignment was the same alignment that was used to generate the reference phylogenetic tree. OTUs that the PyNAST algorithm failed to align were discarded. The PyNAST alignment output was merged with the reference alignment and filtered for gap positions using QIIME (filter_alignment.py) with gap filtering threshold set to 0.99 and entropy threshold set to 0.0001. Identification of Apicomplexans and Chrompodellids reads used a maximum likelihood phylogenetic approach by mapping the OTUs onto our reference tree reference tree using the Evolutionary Placement Algorithm (EPA) of RAxML [67]. OTUs that were not placed within the Apicomplexans and Chrompodellids were removed. Trees using the remaining sequences were built consecutively until no more reads were placed outside our two groups of interest. OTUs and their clustered sequences were then annotated according to their placement. For novel groups containing only short reads we adopted the same annotation as for the environmental exclusive groups retrieved from GenBank. OTUs that were not placed with any previously defined groups were assigned a new name as outlined above. The annotated OTU table and corresponding Sample metadata (SI3 and SI4) were processed for community analysis using QIIME.

## Acknowledgments

We would like to thanks the Hakai Institute and Tula Foundation. This work was funded by a grant from the Canadian Institutes for Health Research (MOP-42517). JdC was supported by a grant from the Tula Foundation to the Centre for Microbial Biodiversity and Evolution and the Marie Curie International Outgoing Fellowship FP7-PEOPLE-2012-IOF - 331450 CAARL.TH was supported by the SNSF grant (PA00P3 145374).

## Supplementary Material

Supplementary Figure 1. Apicomplexan environment distribution heatmap based on the information associated to the sequences retrieved from HTES studies for different environmental features: redox state, open ocean vs coastal, depth, size fraction and, temperature

Supplementary Table 1. Apicomplexan Reference Database built from 18S rRNA sequences bigger than 500bp retrieved from GenBank.

Supplementary Table 2. Taxonomic summary of the HTES reads at the Genus level.

Supplementary Information 1. Apicomplexan 18S rRNA (sequence length >500 bp) reference tree inferred from a maximum likelihood (RAxML) phylogenetic tree (best tree of 1000 and 1000 bootstrap replicates)

Supplementary Information 2. Apicomplexan 18S rRNA EPA-RAxML tree using the HTES reads as query and the Apicomplexan 18S rRNA reference as backbone.

Supplementary Information 3. Apicomplexans HTES reads OTU table.

Supplementary Information 4. Apicomplexans HTES reads Mapping file.

## References

1. Stentiford GD, Feist SW, Stone DM, Peeler EJ, Bass D. Policy, phylogeny, and the parasite. Trends Parasitol. Elsevier Ltd; 2014;30: 274–281. doi:10.1016/j.pt.2014.04.004

2. Fisher MC, Henk DAD a. D a, Briggs CJ, Brownstein JSJSJS, Madoff LCLC, McCraw SLSLSL, et al. Emerging fungal threats to animal, plant and ecosystem health. Nature. Nature Publishing Group; 2012;484: 186–194. doi:10.1038/nature10947

3. Kuris AM, Hechinger RF, Shaw JC, Whitney KL, Aguirre-Macedo L, Boch C a, et al. Ecosystem energetic implications of parasite and free-living biomass in three estuaries. Nature. 2008;454: 515–518. doi:10.1038/nature06970

4. Preston DL, Mischler JA, Townsend AR, Johnson PTJ. Disease Ecology Meets Ecosystem Science. Ecosystems. Springer US; 2016;19: 737–748. doi:10.1007/s10021-016-9965-2

5. Geisen S, Tveit AT, Clark IM, Richter A, Svenning MM, Bonkowski M, et al. Metatranscriptomic census of active protists in soils. ISME J. Nature Publishing Group; 2015;9: 2178–2190. doi:10.1038/ismej.2015.30

6. Simon M, López-García P, Deschamps P, Moreira D, Restoux G, Bertolino P, et al. Marked seasonality and high spatial variability of protist communities in shallow freshwater systems. ISME J. 2015; 1–13. doi:10.1038/ismej.2015.6

7. de Vargas C, Audic S, Henry N, Decelle J, Mahe F, Logares R, et al. Eukaryotic plankton diversity in the sunlit ocean. Science. 2015;348: 1261605. doi:10.1126/science.1261605

8. Mahé F, de Vargas C, Bass D, Czech L, Stamatakis A, Lara E, et al. Parasites dominate hyperdiverse soil protist communities in Neotropical rainforests. Nat Ecol Evol. Macmillan Publishers Limited, part of Springer Nature.; 2017;1: 050997. doi:10.1038/s41559-017-0091

9. Votýpka J, Modrý D, Oborník M, Šlapeta J, Lukeš J. Apicomplexa. Handbook of the Protists. 2017. pp. 1–58. doi:10.1007/978-3-319-32669-6

10. Köhler S, Delwiche CF, Denny P, Tilney L, Webster P, Wilson R, et al. A Plastid of Probable Green Algal Origin in Apicomplexan Parasites. Science (80-). 1997;275: 1485–1489. doi:10.1126/science.275.5305.1485

11. McFadden GI, Yeh E. The apicoplast: now you see it, now you don’t. Int J Parasitol. 2016; doi:10.1016/j.ijpara.2016.08.005

12. Tenter AM, Heckeroth AR, Weiss LM. Toxoplasma gondii: From animals to humans. Int J Parasitol. 2000;30: 1217–1258. doi:10.1016/S0020-7519(00)00124-7

13. Checkley W, White AC, Jaganath D, Arrowood MJ, Chalmers RM, Chen XM, et al. A review of the global burden, novel diagnostics, therapeutics, and vaccine targets for cryptosporidium. Lancet Infect Dis. 2015;15: 85–94. doi:10.1016/S1473-3099(14)70772-8

14. Keeling PJ, Rayner JC. The origins of malaria: there are more things in heaven and earth. Parasitology. 2014; 1–10. doi:10.1017/S0031182014000766

15. Leander BS. Marine gregarines: evolutionary prelude to the apicomplexan radiation? Trends Parasitol. 2008;24: 60–7. doi:10.1016/j.pt.2007.11.005

16. Chambouvet A, Valigurová A, Mesquita L, Richards TA, Jirků M. Nematopsis temporariae (Gregarinasina, Apicomplexa, Alveolata) intracellular infectious agent of tadpole livers. Environ Microbiol Rep. 2016;1–14.

17. Woo YH, Ansari H, Otto TD, Klinger CM, Kolisko M, Michálek J, et al. Chromerid genomes reveal the evolutionary path from photosynthetic algae to obligate intracellular parasites. Elife. 2015;4: 1–41. doi:10.7554/eLife.06974

18. Zoology I, Federation R, Leander BS, Kuvardina ON, Aleoshin V V., Mylnikov AP, et al. Molecular phylogeny and surface morphology of Colpodella edax (Alveolata): insights into the phagotrophic ancestry of apicomplexans. J Eukaryot Microbiol. 2003;50: 334–40. doi:10.1111/j.1550-7408.2003.tb00145.x

19. Oborník M, Modrý D, Lukeš M, Cernotíková-Stříbrná E, Cihlář J, Tesařová M, et al. Morphology, ultrastructure and life cycle of Vitrella brassicaformis n. sp., n. gen., a novel chromerid from the Great Barrier Reef. Protist. 2012;163: 306–23. doi:10.1016/j.protis.2011.09.001

20. Guillou L, Bachar D, Audic S, Bass D, Berney C, Bittner L, et al. The Protist Ribosomal Reference database (PR2): A catalog of unicellular eukaryote Small Sub-Unit rRNA sequences with curated taxonomy. Nucleic Acids Res. Oxford University Press; 2013;41: 597–604. doi:10.1093/nar/gks1160

21. Quast C, Pruesse E, Yilmaz P, Gerken J, Schweer T, Yarza P, et al. The SILVA ribosomal RNA gene database project: Improved data processing and web-based tools. Nucleic Acids Res. 2013;41: 1–7. doi:10.1093/nar/gks1219

22. Balvočiūtė M, Huson DH. SILVA, RDP, Greengenes, NCBI and OTT — how do these taxonomies compare? BMC Genomics. 2017;18: 114. doi:10.1186/s12864-017-3501-4

23. Cavalier-Smith T. Gregarine site-heterogeneous 18S rDNA trees, revision of gregarine higher classification, and the evolutionary diversification of Sporozoa. Eur J Protistol. Elsevier GmbH; 2014;50: 472–495. doi:10.1016/j.ejop.2014.07.002

24. Janouškovec J, Tikhonenkov D V., Burki F, Howe AT, Kolísko M, Mylnikov AP, et al. Factors mediating plastid dependency and the origins of parasitism in apicomplexans and their close relatives. Proc Natl Acad Sci. 2015;112: 10200–10207. doi:10.1073/pnas.1423790112

25. Yuan C, Keeling PJ, Krause PJ, Horák A. Colpodella spp.-like Parasite Infection in Woman, China. Emerg Infect Dis. 2012;18: 125–127.

26. Fredricks DN, Relman DA. Sequence-based identification of microbial pathogens: a reconsideration of Koch’s postulates. Clin Microbiol Rev. 1996;9:18–33.

27. Bhoora R, Franssen L, Oosthuizen MC, Guthrie AJ, Zweygarth E, Penzhorn BL, et al. Sequence heterogeneity in the 18S rRNA gene within Theileria equi and Babesia caballi from horses in South Africa. Vet Parasitol. 2009;159: 112–120. doi:10.1016/j.vetpar.2008.10.004

28. Levin AE, Krause PJ. Transfusion-transmitted babesiosis. Curr Opin Hematol. 2016;23: 573–580. doi:10.1097/MOH.0000000000000287

29. Barta JR, Ogedengbe JD, Martin DS, Smith TG. Phylogenetic position of the adeleorinid coccidia (myzozoa, Apicomplexa, coccidia, eucoccidiorida, Adeleorina) inferred using 18s rDNA sequences. J Eukaryot Microbiol. 2012;59: 171–180. doi:10.1111/j.1550-7408.2011.00607.x

30. Barta JR. The Dactylosomatidae. Advances in Parasitology 1991.

31. Jirků MM, Modrý D, Šlapeta JR, Koudela B, Lukeš J. The phylogeny of Goussia and Choleoeimeria (apicomplexa; eimeriorina) and the evolution of excystation structures in coccidia. Protist. 2002;153: 379–390. doi:10.1078/14344610260450118

32. Davies AJ. The Biology of Fish Haemogregarines. Adv Parasitol. 1995;36: 117–203. doi:10.1016/S0065-308X(08)60491-1

33. Davies AJ, Ball SJ. The Biology of Fish Coccidia. Adv Parasitol. 1993;32: 294–366.

34. Moreira D, López-García P. Are hydrothermal vents oases for parasitic protists? Trends Parasitol. 2003;19: 556–558. doi:10.1016/j.pt.2003.10.005

35. Janouškovec J, Horák A, Barott KL, Rohwer FL, Keeling PJ. Environmental distribution of coral-associated relatives of apicomplexan parasites. ISME J. 2013;7: 444–447. doi:10.1038/ismej.2012.129

36. Mathur V, del Campo J, Kolisko M, Keeling PJ. Global Diversity and Distribution of Close Relatives of Apicomplexan Parasites. Environ Microbiol. 2018;00: 1–10. doi:10.1111/1462-2920.14134

37. Quillfeldt P, Martínez J, Hennicke J, Ludynia K, Gladbach A, Masello JF, et al. Hemosporidian blood parasites in seabirds - A comparative genetic study of species from Antarctic to tropical habitats. Naturwissenschaften. 2010;97: 809–817. doi:10.1007/s00114-010-0698-3

38. Wilson EO. The Little Things That Run the world (The Importance and Conservation of Invertebrates). Conserv Biol. 1987;1: 344–346. doi:10.1111/j.1523-1739.1987.tb00055.x

39. Jones TC, Overstreet RM, Lotz JM, Frelier PF. Paraophioidina scolecoides n.sp., a new aseptate gregarine from cultured Pacific white shrimp Penaeus vannamei. Dis Aquat Organ. 1994;19: 67–75. doi:10.3354/dao019067

40. Valigurová A, Koudela B. Fine structure of trophozoites of the gregarine Leidyana ephestiae (Apicomplexa: Eugregarinida) parasitic in Ephestia kuehniella larvae (Lepidoptera). Eur J Protistol. 2005;41: 209–218. doi:10.1016/j.ejop.2005.05.005

41. Valigurová A. Sophisticated adaptations of Gregarina cuneata (Apicomplexa) feeding stages for epicellular parasitism. PLoS One. 2012;7. doi:10.1371/journal.pone.0042606

42. Sulaiman I. Infectivity and pathogenicity of Ascogregarina culicis (Eugregarinida: Lecudinidae) to Aedes aegypti (Diptera: Culicidae). J Med Entomol. 1992;29: 1–4. doi:10.1093/jmedent/29.1.1

43. Rueckert S, Simdyanov TG, Aleoshin V V., Leander BS. Identification of a divergent environmental DNA sequence clade using the phylogeny of gregarine parasites (apicomplexa) from crustacean hosts. PLoS One. 2011;6: e18163. doi:10.1371/journal.pone.0018163

44. Théodoridès J. Parasitology of Marine Zooplankton. Adv Mar Biol. 1989;25: 117–177. doi:10.1016/S0065-2881(08)60189-3

45. Humes A. How many copepods? Hydrobiologia. 1994;292/293: 1–7.

46. Plischuk S, Meeus I, Smagghe G, Lange CE. Apicystis bombi (Apicomplexa: Neogregarinorida) parasitizing Apis mellifera and Bombus terrestris (Hymenoptera: Apidae) in Argentina. Environ Microbiol Rep. 2011;3: 565–8. doi:10.1111/j.1758-2229.2011.00261.x

47. Alarcón ME, Huang CGCCG, Tsai YYS, Chen WJ, Dubey AK, Wu WJ. Life Cycle and Morphology of Steinina ctenocephali (Ross 1909) comb. nov. (Eugregarinorida: Actinocephalidae), a Gregarine of Ctenocephalides felis (Siphonaptera: Pulicidae) in Taiwan. Zool Stud. 2011;50: 763–772.

48. Field SG, Michiels NK. Parasitism and growth in the earthworm Lumbricus terrestris: fitness costs of the gregarine parasite Monocystis sp. Parasitology. 2005;130: 397–403. doi:10.1017/S0031182004006663

49. Johnson PTJ, Dobson AP, Lafferty KD, Marcogliese DJ, Memmott J, Orlofske SA, et al. When parasites become prey: Ecological and epidemiological significance of eating parasites. Trends Ecol Evol. Elsevier Ltd; 2010;25: 362–371. doi:10.1016/j.tree.2010.01.005

50. Worden AZ, Follows MJ, Giovannoni SJ, Wilken S, Zimmerman AE, Keeling PJ. Rethinking the marine carbon cycle: Factoring in the multifarious lifestyles of microbes. Science. 2015;347: 1257594–1257594. doi:10.1126/science.1257594

51. Bohmann K, Evans A, Gilbert MTP, Carvalho GR, Creer S, Knapp M, et al. Environmental DNA for wildlife biology and biodiversity monitoring. Trends Ecol Evol. Elsevier Ltd; 2014;29: 358–367. doi:10.1016/j.tree.2014.04.003

52. Pawlowski J, Lejzerowicz F, Apotheloz-Perret-Gentil L, Visco JA, Esling P. Protist metabarcoding and environmental biomonitoring: Time for change. European Journal of Protistology. Elsevier GmbH; 2016. doi:10.1016/j.ejop.2016.02.003

53. Stoeck T, Bass D, Nebel M, Christen R, Jones MDM, Breiner H-W, et al. Multiple marker parallel tag environmental DNA sequencing reveals a highly complex eukaryotic community in marine anoxic water. Mol Ecol. 2010;19: 21–31. doi:10.1111/j.1365-294X.2009.04480.x

54. Heger TJ, Giesbrecht IJWW, Gustavsen J, del Campo J, Kellogg CTE, Hoffman KM, et al. High-throughput environmental sequencing reveals high diversity of litter and moss associated protist communities along a gradient of drainage and tree productivity. Environ Microbiol. 2018; doi:10.1111/1462-2920.14061

55. Amaral-Zettler LA, McCliment EA, Ducklow HW, Huse SM. A Method for Studying Protistan Diversity Using Massively Parallel Sequencing of V9 Hypervariable Regions of Small-Subunit Ribosomal RNA Genes. Langsley G, editor. PLoS One. 2009;4: e6372. doi:10.1371/journal.pone.0006372

56. Edgar RC. Search and clustering orders of magnitude faster than BLAST. Bioinformatics. 2010;26: 2460–2461. doi:10.1093/bioinformatics/btq461

57. Katoh K, Standley DM. MAFFT multiple sequence alignment software version 7: Improvements in performance and usability. Mol Biol Evol. 2013;30: 772–780. doi:10.1093/molbev/mst010

58. Capella-Gutiérrez S, Silla-Martínez JM, Gabaldón T. trimAl: a tool for automated alignment trimming in large-scale phylogenetic analyses. Bioinformatics. 2009;25: 1972–1973. doi:10.1093/bioinformatics/btp348

59. Stamatakis A. RAxML version 8: A tool for phylogenetic analysis and post-analysis of large phylogenies. Bioinformatics. 2014;30: 1312–1313. doi:10.1093/bioinformatics/btu033

60. Camacho C, Coulouris G, Avagyan V, Ma N, Papadopoulos J, Bealer K, et al. BLAST+: architecture and applications. BMC Bioinformatics. 2009;10: 421. doi:10.1186/1471-2105-10-421

61. del Campo J, Massana R. Emerging diversity within Chrysophytes, Choanoflagellates and Bicosoecids based on molecular surveys. Protist. Elsevier GmbH.; 2011;162: 435–448. doi:10.1016/j.protis.2010.10.003

62. Del Campo J, Ruiz-Trillo I. Environmental survey meta-analysis reveals hidden diversity among unicellular opisthokonts. Mol Biol Evol. Oxford University Press; 2013;30: 802–805. doi:10.1093/molbev/mst006

63. Huse SM, Mark Welch DB, Voorhis A, Shipunova A, Morrison HG, Eren AM, et al. VAMPS: a website for visualization and analysis of microbial population structures. BMC Bioinformatics. 2014;15: 41. doi:10.1186/1471-2105-15-41

64. Massana R, Gobet A, Audic S, Bass D, Bittner L, Boutte C, et al. Marine protist diversity in European coastal waters and sediments as revealed by high-throughput sequencing. Environ Microbiol. 2015;17: 4035–4049. doi:10.1111/1462-2920.12955

65. Caporaso JG, Bittinger K, Bushman FD, Desantis TZ, Andersen GL, Knight R. PyNAST: A flexible tool for aligning sequences to a template alignment. Bioinformatics. 2010;26: 266–267. doi:10.1093/bioinformatics/btp636

66. Caporaso JG, Kuczynski J, Stombaugh J, Bittinger K, Bushman FD, Costello EK, et al. QIIME allows analysis of high-throughput community sequencing data. Nat Methods. Nature Publishing Group; 2010;7: 335–336. doi:10.1038/nmeth.f.303

67. Berger SA, Stamatakis A. Aligning short reads to reference alignments and trees. Bioinformatics. 2011;27: 2068–2075. doi:10.1093/bioinformatics/btr320

